# Transcriptional Regulators with Broad Expression in the Zebrafish Spinal Cord

**DOI:** 10.1101/2024.02.14.580357

**Authors:** Samantha J. England, Paul C. Campbell, Santanu Banerjee, Richard L. Bates, Ginny Grieb, William F. Fancher, Katharine E. Lewis

## Abstract

**Background:** The spinal cord is a crucial part of the vertebrate CNS, controlling movements and receiving and processing sensory information from the trunk and limbs. However, there is much we do not know about how this essential organ develops. Here, we describe expression of 21 transcription factors and one transcriptional regulator in zebrafish spinal cord.

**Results:** We analyzed the expression of *aurkb, foxb1a, foxb1b, her8a, homeza, ivns1abpb, mybl2b, myt1a, nr2f1b, onecut1*, *sall1a, sall3a, sall3b, sall4, sox2*, *sox19b, sp8b, tsc22d1, wdhd1*, *zfhx3b, znf804a*, and *znf1032* in wild-type and *MIB E3 ubiquitin protein ligase 1* zebrafish embryos. While all of these genes are broadly expressed in spinal cord, they have distinct expression patterns from one another. Some are predominantly expressed in progenitor domains, and others in subsets of post-mitotic cells. Given the conservation of spinal cord development, and the transcription factors and transcriptional regulators that orchestrate it, we expect that these genes will have similar spinal cord expression patterns in other vertebrates, including mammals and humans.

**Conclusions:** Our data identify 22 different transcriptional regulators that are strong candidates for playing different roles in spinal cord development. For several of these genes, this is the first published description of their spinal cord expression.

## Introduction

The spinal cord is a crucial part of the Central Nervous System (CNS), responsible for controlling movements and receiving and processing sensory information from the trunk and the limbs. Even though the spinal cord is relatively simple compared to the brain, there are still fundamental gaps in our knowledge of how it develops and appropriate neuronal and glial spinal cell types are made, maintained and connected into appropriate neuronal circuitry. Transcription factors, proteins that bind DNA and regulate gene expression, play crucial roles in these processes ^1–3^. In addition, regulatory proteins that modulate the complex organization of chromatin to permit transcription factor activity are also essential for the correct development and differentiation of these cells and tissues. However, we have no idea how many transcription factors and transcriptional regulators are expressed in the spinal cord during development and the temporo-spatial expression patterns of many transcription factors and transcriptional regulators are currently unknown. This significantly impedes our ability to identify the regulatory gene networks that orchestrate spinal cord development. To begin addressing this, as part of a larger gene expression screen to identify transcription factor and transcriptional regulator genes that are expressed in the developing spinal cord ^4,5^ we identified 22 genes that are expressed broadly in the zebrafish embryonic spinal cord. When we started this study, the expression of many of these genes was unknown. Since then, some of them have been studied by other groups, although often in other tissues. In most cases, the published expression data is still limited, particularly in the spinal cord and for some of the genes that we analyze in this paper, there are currently no other published reports of their expression in zebrafish embryos. For example, expression of *znf1032*, has, to our knowledge, never been examined in any vertebrate before. There is also no published zebrafish expression data for *homeza, ivns1abpb, mybl2b, onecut1, tsc22d1* or *wdhd1*, although there are some online photographs available from large scale expression screens ^6–12^.

In this report, we describe the spinal cord expression of all 22 of these genes in zebrafish embryos at key developmental stages, and we also show their expression in the brain and other trunk tissues. In addition, we examine whether these genes are expressed in progenitor and/or post-mitotic spinal cells. We analyze their expression in both whole-mount preparations and lateral cross-sections in wild-type (WT) embryos and *MIB E3 ubiquitin protein ligase 1* (*mib1*, formerly called *mind bomb*) mutants. *mib1* encodes a ubiquitin ligase that is part of the Notch pathway, and in *mib1^ta52b^* mutants the vast majority of spinal cord progenitor cells precociously differentiate as early-forming populations of spinal cord post-mitotic neurons, at the expense of later forming neurons and glia ^13–17^. This enables us to distinguish between progenitor domain expression (which should be lost in *mib1* mutants) and post-mitotic expression (which is often, although not always, expanded into additional cells in *mib1* mutants). In combination with analysing WT trunk cross-sections, this allows us to discriminate between expression in progenitor cells and post-mitotic cells. Interestingly, we find that while all of these genes are broadly expressed in the spinal cord, they have distinct expression patterns from one another. Some are expressed in progenitor cells, others in both progenitor cells and different subsets of post-mitotic cells, and others are predominantly expressed in distinct subtypes of post-mitotic cells. This identification of transcription factor and transcriptional regulator genes with distinct expression patterns in progenitor and/or post-mitotic cells is an essential first step in determining the gene regulatory networks that specify the correct development of spinal cord circuitry and function.

## Results

We have performed several different high-throughput gene expression screens to identify transcription factor genes that are expressed in the developing spinal cord ^e.g. 4,5^. In previous studies, we analyzed transcription factors that are expressed by specific types of spinal cord interneurons ^4,5,18–23^. However, we have also identified a subset of 21 transcription factor genes and 1 transcriptional regulator gene that our initial analyses suggested were broadly expressed in the zebrafish embryonic spinal cord, but are not ubiquitious throughout the embryo. Each of the transcription factor genes encodes a protein containing a verified DNA-binding domain, according to functional classification by InterPro (Table 1; ^24^). In addition, we analysed expression of the putative transcriptional regulator gene, *aurkb* - which is involved in chromatin silencing of differentiated post-mitotic mesenchymal stem cells ^25^. As discussed in the introduction, spinal cord expression of some of these genes has not been reported before, while for most of the others, there are only limited descriptions of their expression in the published literature (see Discussion for more detail).

**Table 1.**
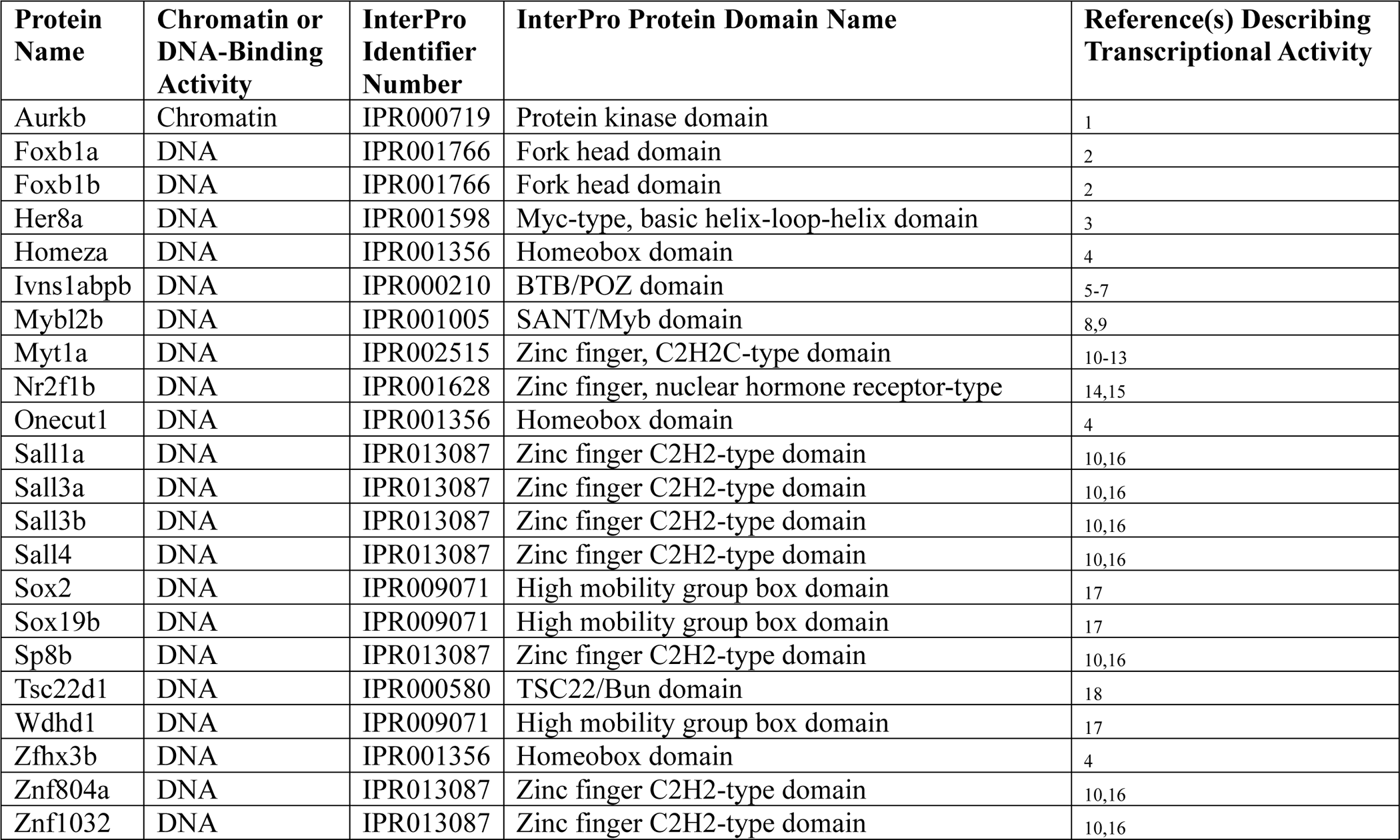
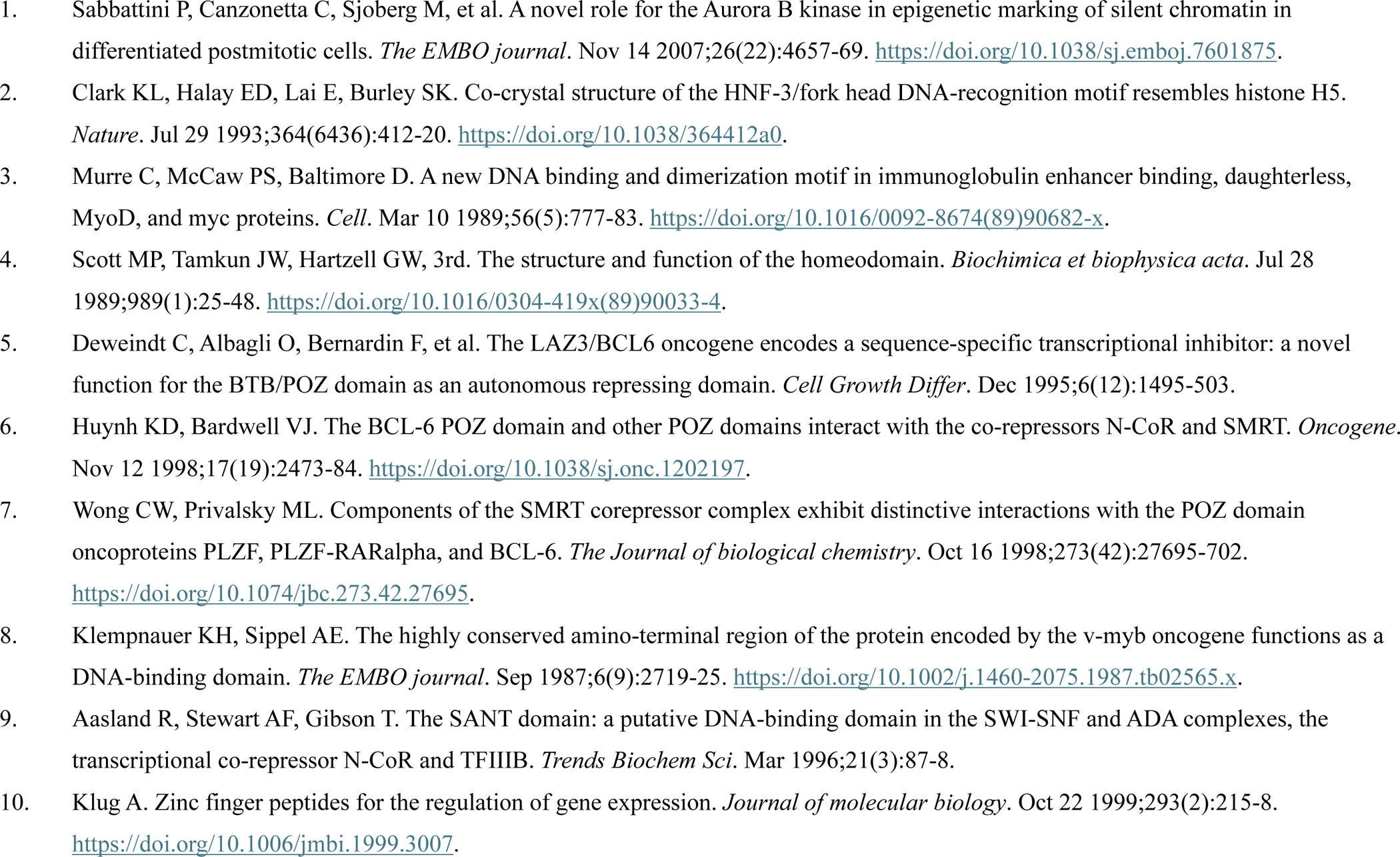
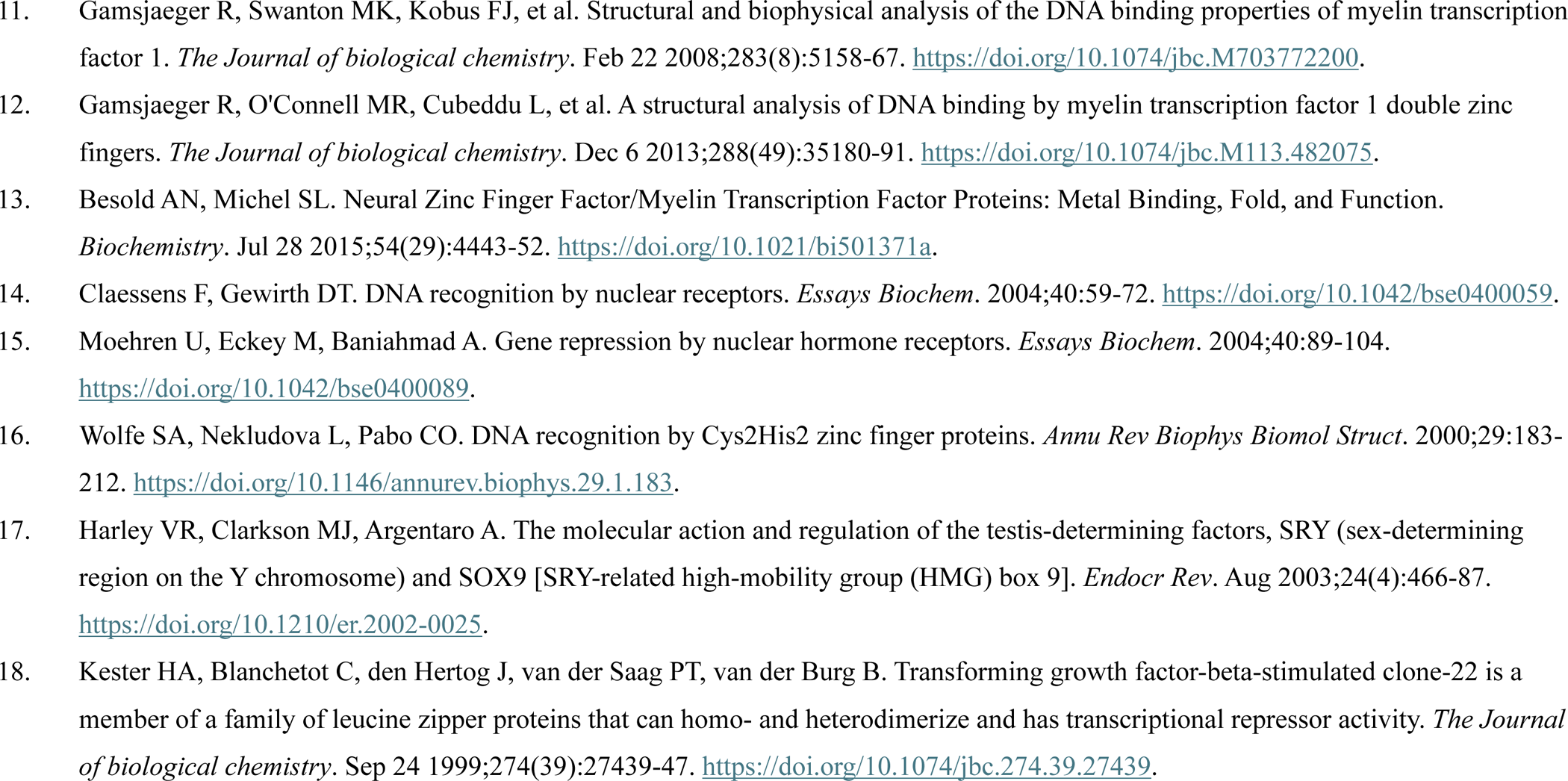
InterPro Identifier Numbers and Domain Names for Transcription Factor and Transcriptional Regulator Proteins Identified in this Study. Column 1 lists proteins analyzed in this study. Column 2 indicates whether the protein has chromatin-remodelling or DNA-binding activity. Columns 3 and 4 contain the InterPro identifier numbers and protein domain names for the transcription-related functional domains contained within each protein ^24^. Column 5 provides references for the papers that document the transcriptional activity of these protein domains and/or the proteins themselves.

As these genes may play important roles in spinal cord development, we analyzed their expression in more detail. At 24 hours post fertilization (h), *aurkb, foxb1a, foxb1b, her8a, homeza, ivns1abpb, mybl2b, myt1a, nr2f1b, onecut1*, *sall1a, sall3a, sall3b, sall4, sox2*, *sox19b, sp8b, tsc22d1, wdhd1*, *zfhx3b, znf804a*, and *znf1032* are all expressed throughout the rostro-caudal and dorso-ventral spinal cord (Fig. 1). All of these genes are also broadly expressed throughout the hindbrain (Figs. 1, 2, 3 and 4, for a schematic of anatomical locations see Fig. 1W) and all of them except *znf804a* and *znf1032* (Fig. 1U, V) have distinct, spatially-restricted expression patterns within the fore- and midbrain. In contrast, *znf804a* and *znf1032* are more broadly expressed in these brain regions (Figs. 1 and 3). In addition, *aurkb*, *ivns1abpb*, *mybl2b*, *sall4*, *wdhd1*, *znf804a*, and *znf1032* are variably expressed in the blood (located in the dorsal aorta and cardinal vein) beneath the notochord (Fig. 1A, F, G, N, S, U, V, W).

**Figure 1.**
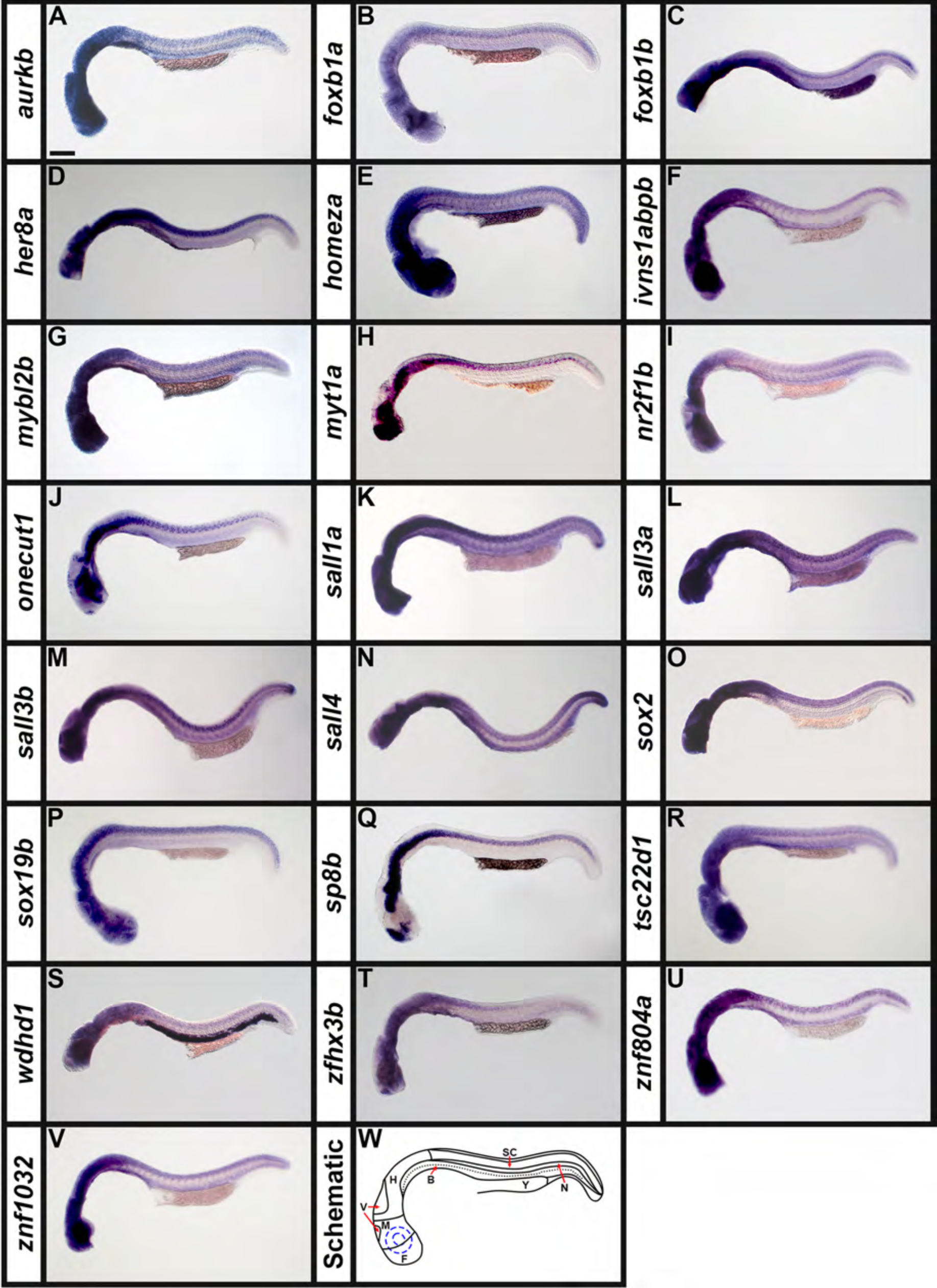
Broad Expression of Transcription Factor and Transcriptional Regulator Genes in Wholemount Zebrafish Embryos at 24 h. (A-V) Lateral views of wholemount WT zebrafish embryos at 24 h. A minimum of 5 embryos were analysed in detail per gene to determine the representative expression pattern (see Experimental Procedures for further details). (W) Schematic of a lateral view of a 24 h wholemount zebrafish embryo. F = forebrain, M = midbrain, H = hindbrain, V = mid- and hindbrain ventricles, SC = spinal cord, N = notochord, B = blood (black dotted line indicates boundary between dorsal aorta and cardinal vein), Y = yolk and the eye and lens are indicated with concentric blue dotted circles. In all panels, rostral is left and dorsal is up. Transcriptional regulator gene (A) *aurkb,* and transcription factor genes (B) *foxb1a,* (C) *foxb1b,* (D) *her8a,* (E) *homeza,* (F) *ivns1abpb,* (G) *mybl2b,* (H) *myt1a,* (I) *nr2f1b,* (J) *onecut1*, (K) *sall1a,* (L) *sall3a,* (M) *sall3b,* (N) *sall4,* (O) *sox2* (P), *sox19b,* (Q) *sp8b,* (R) *tsc22d1,* (S) *wdhd1*, (T) *zfhx3b,* (U) *znf804a*, and (V) *znf1032* are all broadly expressed throughout the rostro-caudal and dorso-ventral spinal cord and all 22 of these genes are also variably expressed in the brain. (A) *aurkb*, (F) *ivns1abpb*, (G) *mybl2b*, (N) *sall4*, (S) *wdhd1*, (U) *znf804a*, and (V) *znf1032* are also variably expressed in the blood beneath the notochord. (N) *sall4 in situ* hybridization experiments were performed with the molecular crowding reagent Dextran Sulfate (see Experimental Procedures for rationale). All other *in situ* hybridization experiments in this figure were performed without this reagent. Scale bar: 200 µm.

**Figure 2.**
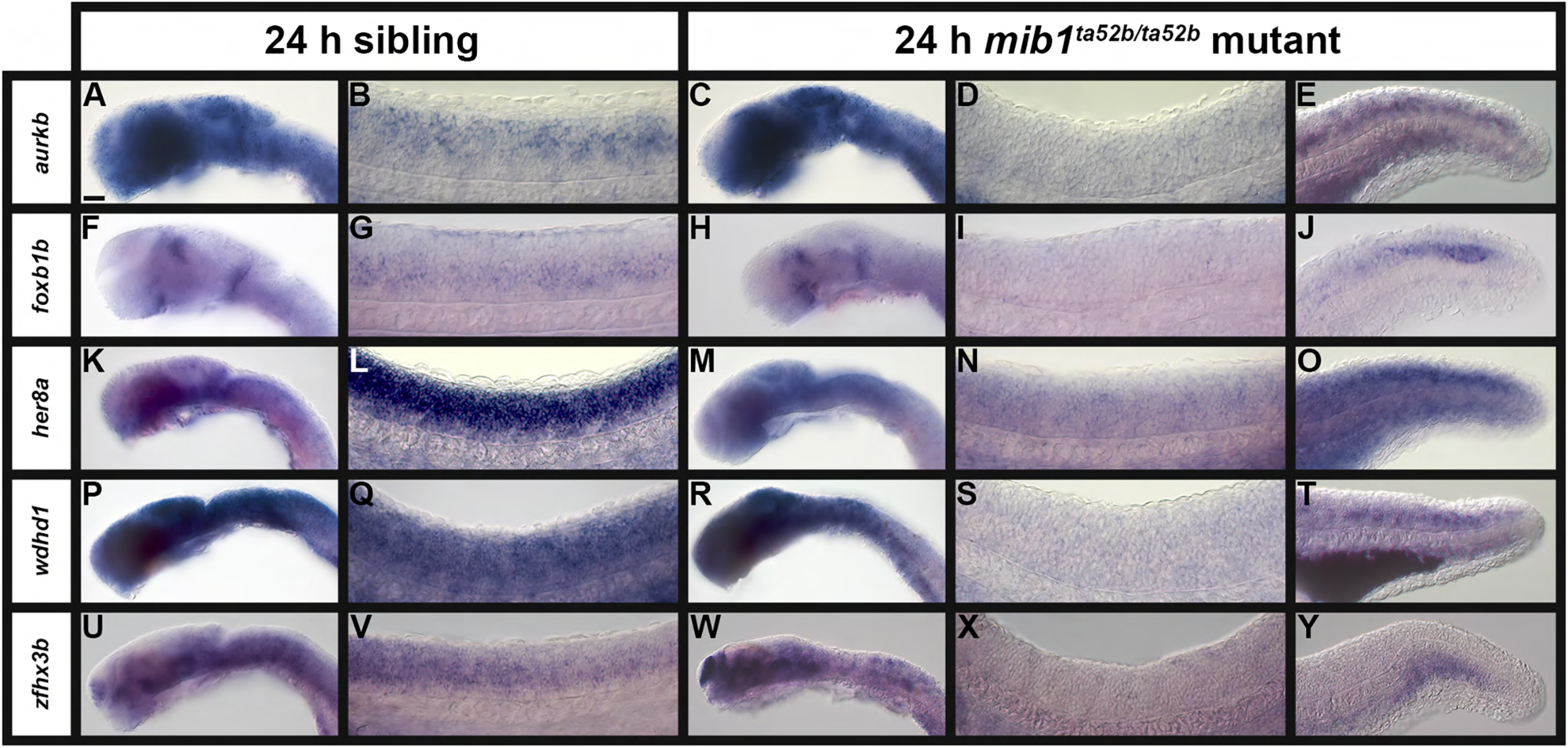
A Subset of Transcription Factor and Transcriptional Regulator Genes Lose Expression in the Spinal Cord of Zebrafish *mib1^ta52b^* Mutant Embryos at 24 h. (A-Y) Lateral views of (A, C, F, H, K, M, P, R, U, W) head, (B, D, G, I, L, N, Q, S, V, X) spinal cord, and (E, J, O, T, Y) tail in (A, B, F, G, K, L, P, Q, U, V) sibling and (C-E, H-J, M-O, R-T, W-Y) *mib1^ta52b^* mutant embryos at 24 h. Rostral, left. Dorsal, up. A minimum of 5 embryos were analysed per gene for each genotype to determine representative expression patterns (see Experimental Procedures). None of the *in situ* hybridization experiments in this figure were performed with the molecular crowding reagent Dextran Sulfate. Scale bar: (A, C, F, H, K, M, P, R, U, W) 50 µm, (B, D, G, I, L, N, Q, S, V, X) 20 µm, and (E, J, O, T, Y) 25 µm.

**Figure 3.**
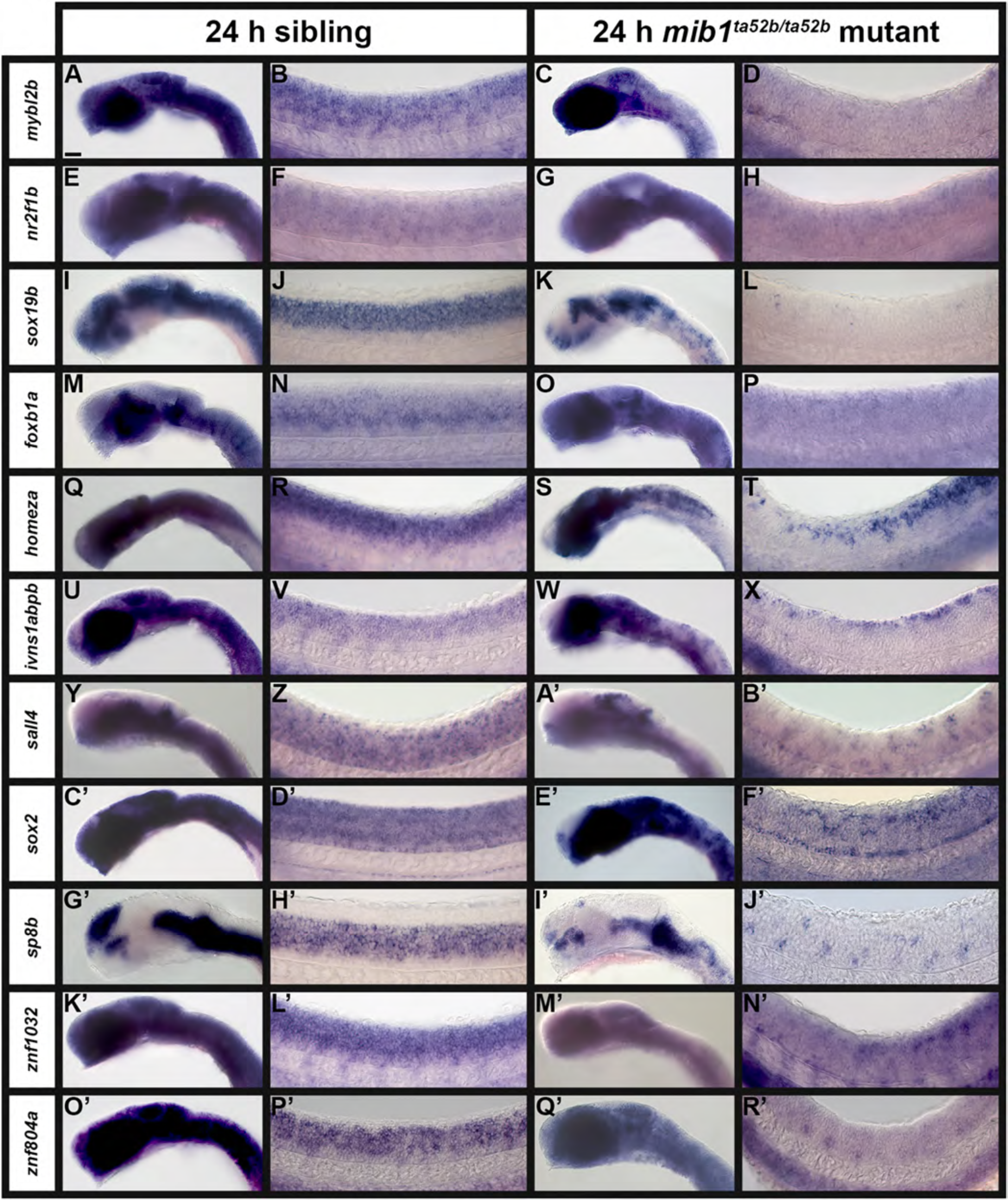
A Subset of Transcription Factor Genes Show Reduced Expression in the Spinal Cord of Zebrafish *mib1^ta52b^* Mutant Embryos at 24 h. Lateral views of (A, C, E, G, I, K, M, O, Q, S, U, W, Y, A’, C’, E’, G’, I’, K’, M’, O’, Q’) head, and (B, D, F, H, J, L, N, P, R, T, V, X, Z, B’, D’, F’, H’, J’, L’, N’, P’, R’) spinal cord in (A, B, E, F, I, J, M, N, Q, R, U, V, Y, Z, C’, D’, G’, H’, K’, L’, O’, P’) sibling and (C, D, G, H, K, L, O, P, S, T, W, X, A’, B’, E’, F’, I’, J’, M’, N’, Q’, R’) *mib1^ta52b^* mutant embryos at 24 h. Rostral, left. Dorsal, up. A minimum of 5 embryos were analysed per gene for each genotype to determine representative expression patterns (see Experimental Procedures). (Y-B’) *sall4 in situ* hybridization experiments were performed with the molecular crowding reagent Dextran Sulfate (see Experimental Procedures for rationale). All other *in situ* hybridization experiments in this figure were performed without this reagent. Scale bar: (A, C, E, G, I, K, M, O, Q, S, U, W, Y, A’, C’, E’, G’, I’, K’, M’, O’, Q’) 50 µm, (B, D, F, H, J, L, N, P, R, T, V, X, Z, B’, D’, F’, H’, J’, L’, N’, P’, R’) 20 µm.

**Figure 4.**
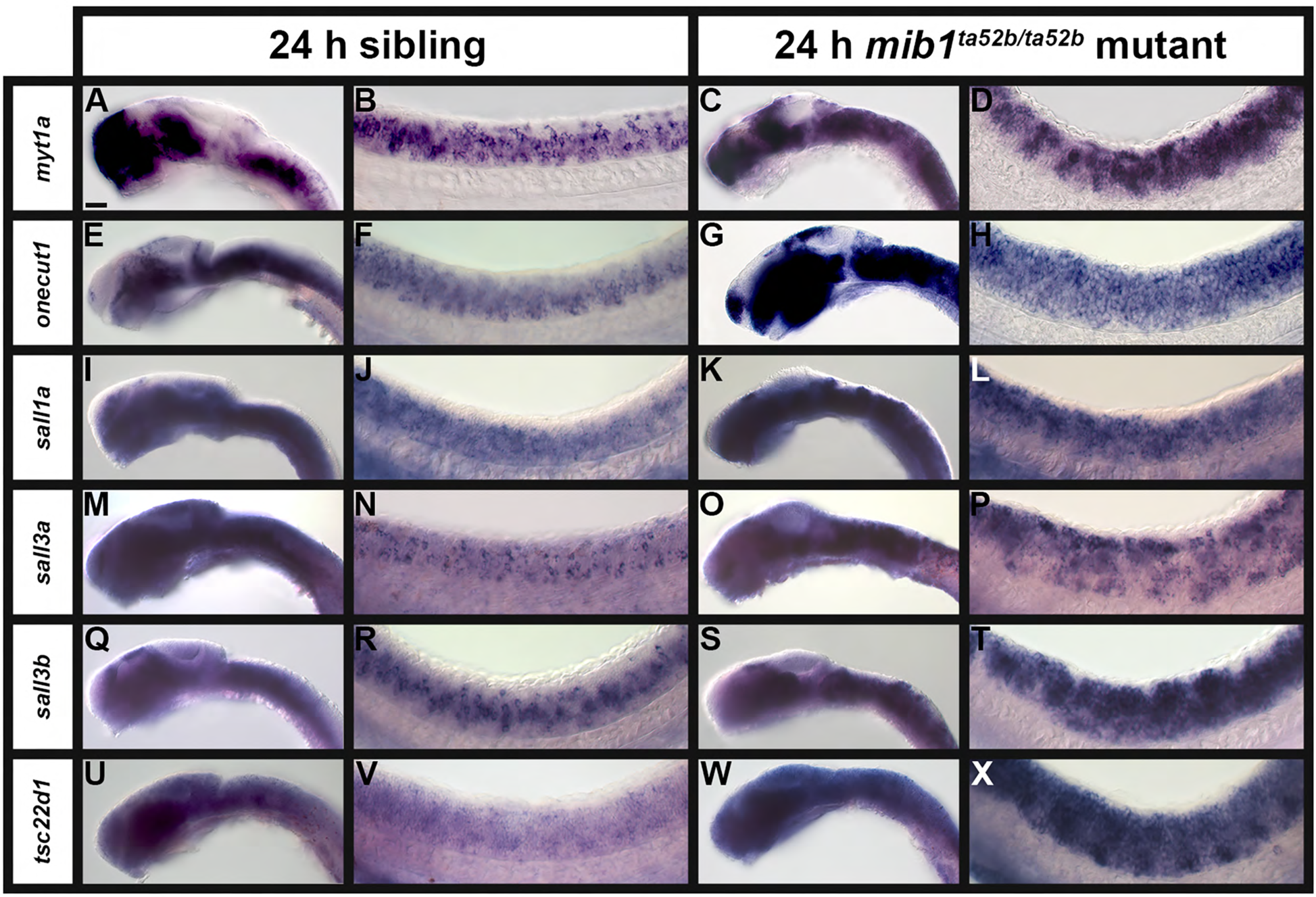
A Subset of Transcription Factor Genes Show Expanded Expression in the Spinal Cord of Zebrafish *mib1^ta52b^* Mutant Embryos at 24 h. Lateral views of (A, C, E, G, I, K, M, O, Q, S, U, W) head, and (B, D, F, H, J, L, N, P, R, T, V, X) spinal cord in (A, B, E, F, I, J, M, N, Q, R, U, V) sibling and (C, D, G, H, K, L, O, P, S, T, W, X) *mib1^ta52b^* mutant embryos at 24 h. Rostral, left. Dorsal, up. A minimum of 5 embryos were analysed per gene for each genotype to determine representative expression patterns (see Experimental Procedures). In most of these cases, gene expression can be observed in additional spinal cord regions in *mib1^ta52b^* mutants compared to WT and sibling embryos, because of the expansion of specific populations of post-mitotic cells into neighboring locations along both dorsal-ventral and medial-lateral axes (see Figure 8). None of the *in situ* hybridization experiments in this figure were performed with the molecular crowding reagent Dextran Sulfate. Scale bar: (A, C, E, G, I, K, M, O, Q, S, U, W) 50 µm, (B, D, F, H, J, L, N, P, R, T, V, X) 20 µm.

The spinal cord expression of *aurkb, foxb1a, homeza, myt1a, onecut1, sall1a, sall3b, sall4, sox2, sox19b, sp8b*, *zfhx3b*, and *znf1032* persists at 36 h (Fig. 5A, B, E, H, J, K, M-Q, T, V). In contrast, the spinal cord expression of *foxb1b, her8a, ivns1abpb, mybl2b, nr2f1b, sall3a, tsc22d1, wdhd1*, and *znf804a* decreases by this stage, either persisting most strongly in ventral spinal cord (*foxb1b*, *her8a*, *ivns1abpb*, *mybl2b*, *sall3a*, and *znf804a*), only very weakly in the spinal cord (*nr2f1b* and *tsc22d1*), or only in very few spinal cells (*wdhd1*) (Fig. 5C, D, F, G, I, L, R, S, U). *her8a*,*sox2* and *znf804a* are now also expressed in neuromasts, deposited at intervals along the length of the embryo by the migrating lateral line primordium (black arrows, Fig. 5W, X, E’). The variable expression of *aurkb, ivns1abpb, mybl2b, sall4, wdhd1, znf804a* and *znf1032* in the blood (dorsal aorta and cardinal vein immediately beneath the notochord) persists at this stage (Fig. 5Z-F’). Interestingly, *wdhd1* is expressed in individual blood cells at this stage (Fig. 5D’).

**Figure 5.**
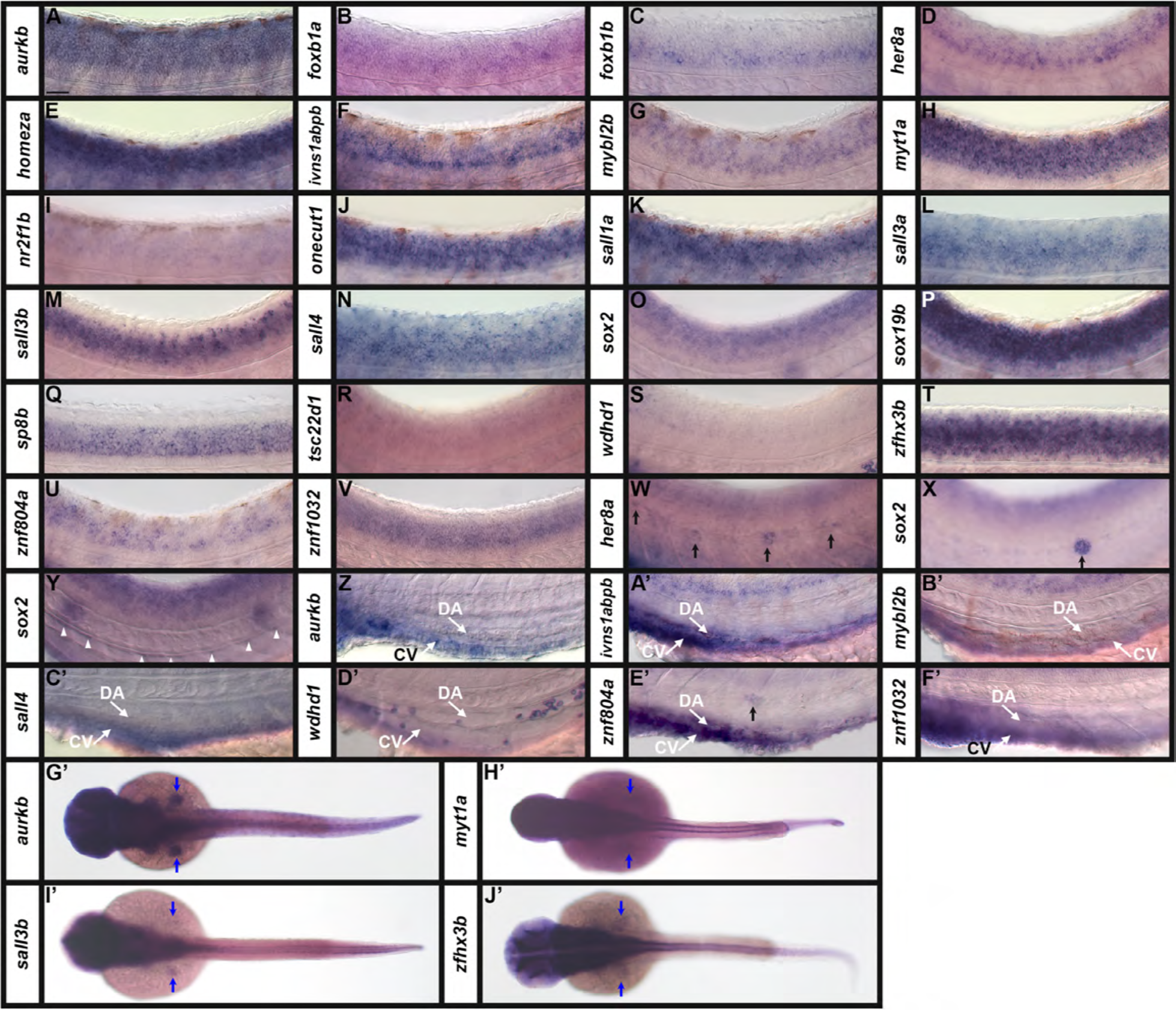
Transcription Factor and Transcriptional Regulator Gene Expression in Zebrafish Spinal Cord and Mesendodermal Tissues at 36 h. Lateral (A-F’) views of spinal cord (A-V), neuromasts of the lateral line primordium (W, X, E’), the hypochord (Y), and the blood (Z-F’) in WT zebrafish embryos at 36 h. Rostral, left. Dorsal, up. (G’-J’) Dorsal wholemount views of WT zebrafish embryos at 36 h. Rostral, left. A minimum of 5 embryos were analysed per gene to determine the representative expression pattern (see Experimental Procedures). (A-V) In the spinal cord views, the morphological boundary between the ventral spinal cord and the notochord is visible towards the bottom of the panel and the spinal cord is in focus. In panels (W, X, E’) the focal plane is more lateral, and the somites and lateral line primordium are in focus. Black arrows (W, X, E’) indicate neuromasts, deposited at intervals along the length of the embryo by the migrating lateral line primordium. White arrowheads (Y) indicate the hypochord, at the ventral interface between the notochord and the blood. (Z-F’) Expression in the blood is observed in the dorsal aorta (DA, white arrow) and cardinal vein (CV, white arrow), beneath the notochord. (G’-J’) Expression in the pectoral fin buds is indicated (blue arrows). *in situ* hybridization experiments with (L) *sall3a,* (M, I’) *sall3b,* (N, C’) *sall4*, and (T, J’) *zfhx3b* were performed with the molecular crowding reagent Dextran Sulfate (see Experimental Procedures for rationale). All other *in situ* hybridization experiments in this figure were performed without this reagent. Scale bar: (A-F’) 30 µm, (G’-J’) 130 µm.

While we concentrated predominantly on spinal cord expression of these genes, at 36 h we also, as at 24 h, observed different brain expression patterns (Figure 6, for a schematic of anatomical locations within the brain, see Fig. 6W). *myt1a, sox19b, sp8b* and *zfhx3b* have distinct expression patterns in the pallium and subpallium of the telencephalon (dorsal forebrain) (Fig. 6H, P, Q, T, W). In the diencephalon (ventral forebrain), *homeza, onecut1, sox2* and *wdhd1* are expressed in the habenula, and there is variable expression of *nr2f1b, onecut1, sox19b, sp8b, wdhd1* and *zfhx3b* in the preoptic region, *sox19b* and *myt1a* in the thalamic region, and *aurkb, foxb1a, foxb1b, myt1a, wdhd1* and *znf1032* in the hypothalamus (Fig. 6A, B, C, E, H, I, J, O, P, Q, S, T, V, W). We also observed distinct expression patterns within the hindbrain. For example, *aurkb, foxb1a* and *foxb1b* are expressed in both the tectum and tegmentum, at the boundary with the hindbrain (Fig. 6A, B, C, W). In contrast, *ivns1abpb, mybl2b, myt1a, nr2f1b, sox2, tsc22d1, wdhd1* and *zfhx3b* are variably expressed in the tectum (dorsal midbrain) (Fig. 6F, G, H, I, O, R, S, T, W). Within the tegmentum (ventral midbrain), *myt1a, nr2f1b, sox2* and *zfhx3b* are more broadly expressed than they are in the tectum, whilst *wdhd1* expression is more spatially defined in the tegmentum compared to the tectum (Fig. 6H, I, O, T, W). We also observed expression of *onecut1, sox19b, sp8b* and *znf1032* in the tegmentum (Fig. 6J, P, Q, R, W). Within the hindbrain (cerebellum), *myt1a, nr2f1b, sox2, sox19b, sp8b, zfhx3b* and *znf1032* are widely expressed throughout the structure. In contrast, *her8a, homeza, ivns1abpb, mybl2b* and *wdhd1* are restricted to expression in the rhombomeres (white arrowheads) (Fig. 6D, E, F, G, H, I, O, P, Q, S, T, V, W). In comparison to the genes described above, we observed extensive expression of *sall1a, sall3a, sall3b, sall4, tsc22d1* and *znf804a* throughout all brain regions (Fig. 6K, L, M, N, R, U, W). Lateral to the brain, we also observed expression of *aurkb* and *tsc22d1* in the otic vesicles (white asterisks, Fig. 6A, R), *aurkb, onecut1, sall3b* and *sox2* in the branchial (gill) mesenchyme adjacent to the hindbrain (black arrows) (Fig. 6A, J, M, O), and *onecut1* and *sp8b* in the olfactory bulbs (blue arrowheads) (Fig. 6J and data not shown).

**Figure 6.**
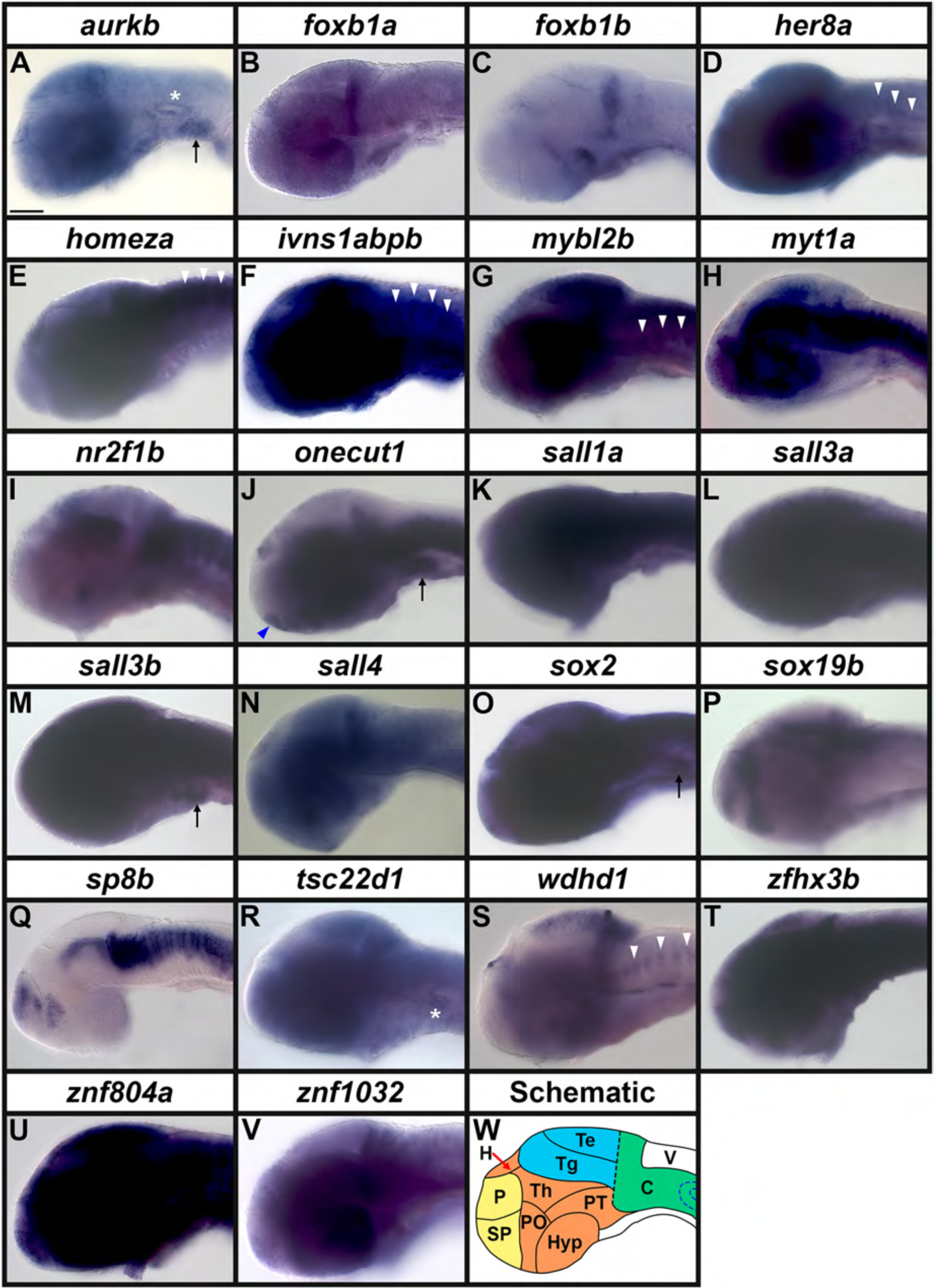
Transcription Factor and Transcriptional Regulator Gene Expression in Zebrafish Brain at 36 h. (A-V) Lateral views of heads in WT zebrafish embryos at 36 h. A minimum of 5 embryos were analysed per gene (see Experimental Procedures for further details). (W) Schematic of a lateral view of the head of a 36 h zebrafish embryo. The telencephalon (dorsal forebrain, yellow) consists dorsally and ventrally of the pallium (P) and subpallium (SP) respectively. The diencephalon (ventral forebrain, orange) consists of the habenula (H), the hypothalamus (Hyp), the Thalamic region (Th), the Posterior Tuberculum (PT) and the preoptic region (PO). The midbrain (blue) consists dorsally and ventrally of the tectum (Te) and the tegmentum (Tg) resepectively. The hindbrain (green) consists of the cerebellum (C). V = ventricle in the hindbrain. Black dotted line = midbrain-hindbrain boundary. Blue dotted lines = otic vesicle. Transcriptional regulator gene (A) *aurkb,* and transcription factor genes (B) *foxb1a,* (C) *foxb1b,* (D) *her8a,* (E) *homeza,* (F) *ivns1abpb,* (G) *mybl2b,* (H) *myt1a,* (I) *nr2f1b,* (J) *onecut1*, (K) *sall1a,* (L) *sall3a,* (M) *sall3b,* (N) *sall4,* (O) *sox2* (P), *sox19b,* (Q) *sp8b,* (R) *tsc22d1,* (S) *wdhd1*, (T) *zfhx3b,* (U) *znf804a*, and (V) *znf1032* are all all variably expressed in the brain. (D, E, F, G, S) White arrowheads depict weak expression in the rhomobomeres of the hindbrain. (A, R) *aurkb* and *tsc22d1* are expressed in the otic vesicles (white asterisks). (A, J, M, O) *aurkb, onecut1, sall3b* and *sox2* are expressed in the branchial (gill) mesenchyme (black arrow). (J and data not shown) *onecut1* and *sp8b* are expressed in the olfactory bulbs (blue arrowhead). (N) *sall4 in situ* hybridization experiments were performed with the molecular crowding reagent Dextran Sulfate (see Experimental Procedures for rationale). All other *in situ* hybridization experiments in this figure were performed without this reagent. Scale bar: 100 µm.

To determine whether each of these genes is expressed in spinal cord progenitor cells and/or post-mitotic cells we analyzed their expression in spinal cord cross-sections and *mib* mutants at 24 h. Our WT spinal cord cross-sections show that most of these genes, *aurkb, foxb1a, foxb1b*, *her8a, homeza*, *ivns1abpb, mybl2b, nr2f1b, sall1a, sall3b*, *sall4, sox2, sox19b, sp8b, tsc22d1, wdhd1, zfhx3b, znf804a,* and *znf1032*, are expressed in both the medial (progenitor), and lateral (post-mitotic) domains of the spinal cord (Fig. 7). These genes are also expressed throughout the dorso-ventral axis of the spinal cord, except for *sall3b* and *sp8b*, which are only expressed in the ventral two-thirds of the spinal cord (Fig. 7M, Q), and *foxb1b* and *wdhd1*, which are expressed throughout the dorso-ventral medial (progenitor) domains of the spinal cord, but are only expressed in the ventral two-thirds of the lateral (post-mitotic) domains of the spinal cord (Fig. 7C, S). In contrast, *myt1a* and *onecut1* are only expressed in the lateral (post-mitotic) domains of the spinal cord (Fig. 7H, J), and if *sall3a* is expressed in the medial spinal cord, that expression is much weaker than the lateral expression (Fig. 7L). *myt1a* and *onecut1* expression extends to all dorso-ventral domains of the spinal cord, but *sall3a* expression is absent in the dorsal-most spinal cord (Fig. 7H, J, L). Consistent with the whole-mount analyses (Fig. 1), *aurkb*, *ivns1abpb*, *mybl2b*, *sall4*, *wdhd1*, *znf804a*, and *znf1032* expression is also visible in the blood immediately beneath the notochord (Fig. 7A, F, G, N, S, U, V, W). In addition, *sox2* is also expressed in the hypochord, at the ventral interface between the notochord and the blood (Fig. 7O, for a schematic of anatomical locations see Fig. 7W) and *tsc22d1* is expressed in the pronephros, which are tubular structures paired ventrally, lateral to the blood, on each side of the cross-section (Fig. 7R, and see Fig. 7W).

**Figure 7.**
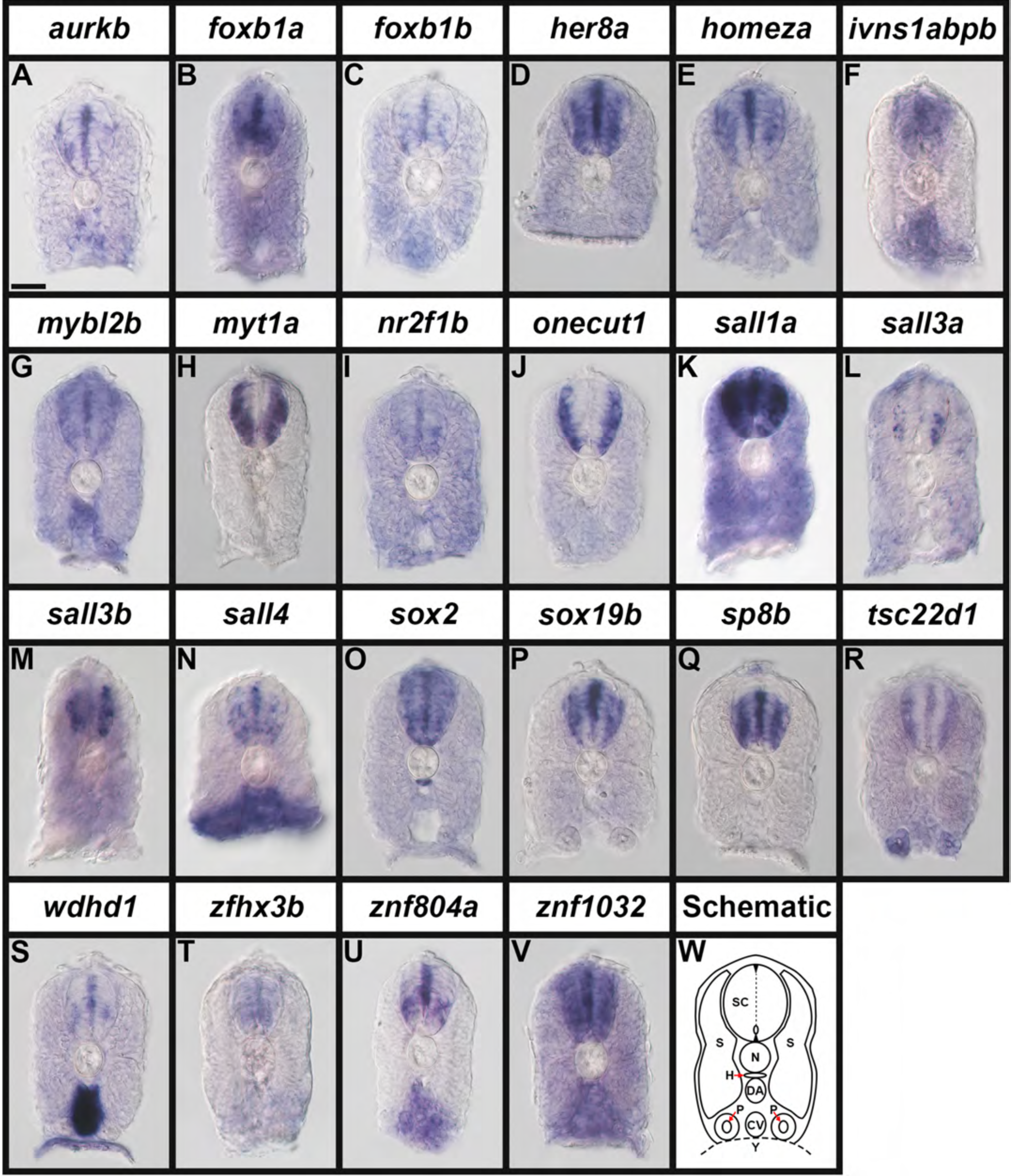
Broad Expression of Transcription Factor and Transcriptional Regulator Genes in Zebrafish Spinal Cord at 24 h. (A-V) Cross-section views of trunk expression of transcriptional regulator gene (A) *aurkb,* and transcription factor genes (B) *foxb1a,* (C) *foxb1b*, (D) *her8a,* (E) *homeza*, (F) *ivns1abpb,* (G) *mybl2b,* (H) *myt1a,* (I) *nr2f1b,* (J) *onecut1,* (K) *sall1a,* (L) *sall3a,* (M) *sall3b*, (N) *sall4,* (O) *sox2,* (P) *sox19b,* (Q) *sp8b,* (R) *tsc22d1,* (S) *wdhd1,* (T) *zfhx3b,* (U) *znf804a,* and (V) *znf1032* in WT zebrafish embryos at 24 h. Dorsal, up. A minimum of 5 embryos were analysed per gene to determine the representative expression pattern (see Experimental Procedures). As indicated in the schematic cross-section (W), the spinal cord (SC), is located above the notochord (N), which is above the hypochord (H, indicated with red arrow), dorsal aorta (DA), and the cardinal vein (CV). The somites (S) can be seen on both sides of these tissues and the pronephros tubes (P, indicated with red arrows) are ventral, either side of the cardinal vein. Within the spinal cord, the dotted line indicates the midline, the small oval indicates the central canal and the small black triangles indicate the roof plate and floor plate. (N) *sall4 in situ* hybridization experiments were performed with the molecular crowding reagent Dextran Sulfate (see Experimental Procedures for rationale). All other *in situ* hybridization experiments in this figure were performed without this reagent. Scale bar: 30 µm.

To further confirm whether these genes are expressed in progenitor cells or post-mitotic cells, we also examined their expression in 24 h *mib1^ta52b^* mutants. As described in the introduction, the vast majority of spinal cord progenitor cells precociously differentiate as early-forming populations of post-mitotic neurons in these mutants, at the expense of later forming neurons and glia ^13–17^. Therefore, spinal cord progenitor domain expression is lost, and post-mitotic expression is usually expanded to additional cells, in these mutants (e.g. see Fig. 8A-D).

**Figure 8.**
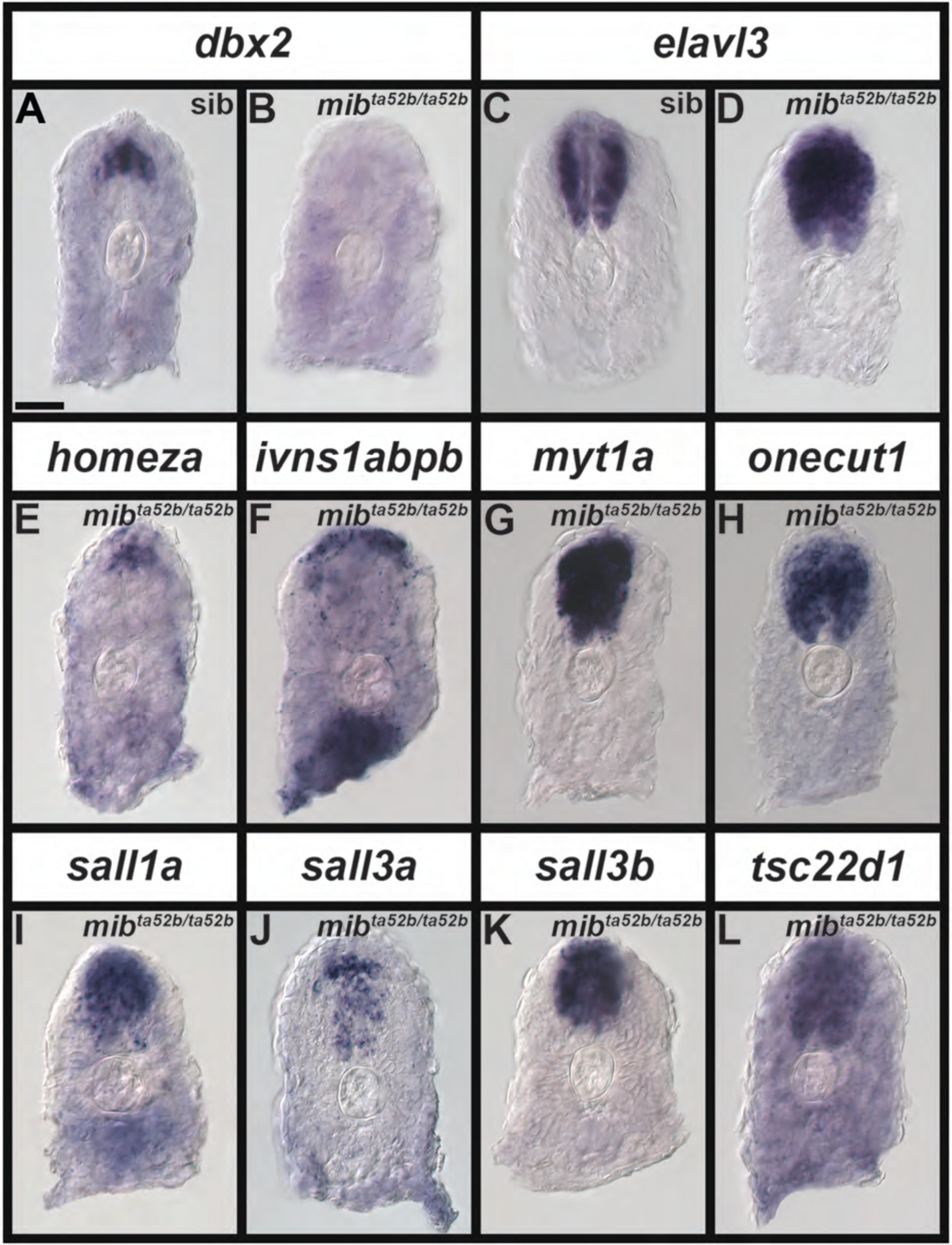
Expression of a Subset of Transcription Factor Genes is Either Lost in Progenitor Domains and/or Expanded in Post-Mitotic Domains in the Spinal Cord of Zebrafish *mib1^ta52b^* Mutant Embryos at 24 h. (A-L) Cross-section views of trunk expression of transcription factor genes (A, B) *dbx2* (expressed in a spinal cord progenitor domain), (C-D) *elavl3* (expressed by all spinal cord post-mitotic cells), (E) *homeza*, (F) *ivns1abpb,* (G) *myt1a*, (H) *onecut1,* (I) *sall1a,* (J) *sall3a,* (K) *sall3b*, and (L) *tsc22d1* in (A, C) sibling and (B, D, E, F, G, H, I, J, K, L) *mib1^ta52b^* mutant embryos at 24 h. Dorsal, up. A minimum of 5 embryos were analysed per gene per genotype to determine representative expression patterns (see Experimental Procedures). For schematic, please see Fig. 7W. The expression patterns of *homeza, ivns1abpb, myt1a, onecut1, sall1a, sall3a, sall3b* and *tsc22d1* in sibling spinal cords at 24 h (data not shown) is identical to that in 24 h WT embryos shown in Fig. 7. None of the *in situ* hybridization experiments in this figure were performed with the molecular crowding reagent Dextran Sulfate. Scale bar: 30 µm.

*mib1^ta52b^* mutant embryos and their siblings were processed and stained identically and we analyzed at least 5 embryos per gene per genotype (see Experimental Procedures). We found that expression of five of the genes, *aurkb, foxb1b, her8a, wdhd1* and *zfhx3b,* is lost from the spinal cord in all but the very caudal tail in *mib1^ta52b^* mutants (Fig. 2), suggesting that these genes are all expressed in spinal cord progenitor cells. Interestingly though, brain expression of these genes is largely unchanged in these mutants, with the exception that *her8a* expression is slightly less intense, and *zfhx3b* expression is slightly expanded in the telencephalon (Fig. 2M, W). *aurkb* and *wdhd* expression in the blood is also unaffected (Fig. 2E, T).

Eleven other genes also lose a lot of their normal spinal cord expression in *mib1^ta52b^* mutants, but in contrast to the five genes discussed above, they are still expressed in distinct subsets of spinal cord cells (Fig. 3). Expression of *mybl2b, nr2f1b, sox19b*, and *foxb1a* is lost in all but a few dorsal spinal cord cells, whereas expression of *homeza* and *ivns1abpb* persists in more cells but only in dorso-caudal spinal cord (Fig. 3D, H, L, P, T, X). In contrast, small clusters of cells in both dorsal and ventral spinal cord still express *sall4, sox2*, *sp8b*, and *znf1032*, whereas *znf804a* is only expressed in small clusters of ventral spinal cord cells (Fig. 3B’, F’, J’, N’, R’). These data suggest that these genes are expressed broadly in spinal cord progenitor cells, and also in a small number of post-mitotic spinal cord cells. Interestingly, and in contrast to the five genes discussed above (Fig. 2), all eleven of these genes also have a dramatic reduction in hindbrain expression in *mib1^ta52b^* mutants. However, consistent with the five genes discussed above, their expression in the fore- and midbrain regions is largely unchanged in *mib1^ta52b^* mutants, with the exception of *sall4*, which is expanded into a slightly larger territory of the dorsal midbrain, and *sox2* and *sp8b*, which are almost lost in the dorsal forebrain (Fig. 3A’, E’, I’). Expression of *mybl2b*, *ivns1abpb*, *sall4*, *znf1032*, and *znf804a* in the blood is unaffected in *mib1^ta52b^* mutants (Fig. 3D, X, B’, N’, R’).

Unlike the 16 genes discussed above, expression of the remaining six genes, *myt1a, onecut1, sall1a, sall3a, sall3b*, and *tsc22d1*, persists and is expanded into additional cells in the spinal cord of *mib1^ta52b^* mutant embryos (Fig. 4). This suggests that these genes are expressed by post-mitotic spinal cord cells. *myt1a*, *onecut1*, *sall1a*, and *sall3a* expression is also expanded to include additional cells throughout the brain in *mib1^ta52b^* mutants, including the hindbrain (Fig. 4C, G, K, O). In contrast, *sall3b* and *tsc22d1* expression is only expanded into additional cells in the hindbrain and largely unchanged elsewhere in the brain (Fig. 4S, W).

We also performed cross-sectional analyses for some of the genes that still had spinal cord expression in *mib1^ta52b^* mutants (Fig. 8). As a comparison, we first analyzed the expression of a known spinal cord progenitor domain marker, *dbx2*, and post-mitotic marker, *elavl3,* in spinal cord cross-sections (Fig. 8A-D). In WT zebrafish embryos, *dbx2* is expressed in the spinal cord progenitor domains pd5, pd6, p0 and p1 at 24 h ^26,27^. Our cross-sections show that this progenitor domain expression occupies a medial mid-dorsoventral position in the spinal cord of sibling embryos but is lost completely in the spinal cord of *mib1^ta52b^* mutants (Fig. 8A, B). In contrast, *elavl3* is expressed by all post-mitotic neurons in the zebrafish spinal cord ^28–30^. In sibling embryo cross sections, *elavl3* is expressed by post-mitotic cells in the lateral spinal cord. It is not expressed by progenitor cells in the medial spinal cord. In *mib1^ta52b^* mutant cross-sections, *elavl3* expression persists laterally but also expands medially to include every cell in the spinal cord (Fig. 8C, D), consistent with spinal progenitor cells precociously differentiating and becoming post-mitotic in these mutants.

The spinal cord expression of *homeza* and *ivns1abpb* in *mib1^ta52b^* mutants appears, in lateral view, to be in specific post-mitotic dorsal spinal cord domains. To further confirm this we analyzed spinal cord cross sections for these two genes. Consistent with the lateral views, the only remaining expression of *homeza* and *ivns1abpb* is in the very dorsal spinal cord, where it extends across the medio-lateral axis, as we would expect for a specific population of post-mitotic cells (Fig. 8E, F). This is in contrast to the expression of these genes in WT and sibling embryos, where they are expressed in both medial progenitor, and lateral post-mitotic cells along most of the dorsoventral axis of the spinal cord (Fig. 7E, F and data not shown). This suggests that all progenitor domain expression of these genes has been lost and that post-mitotic expression only persists in the dorsal spinal cord. We also performed cross-sectional analyzes for some of the genes that, from the lateral views, appear to be expressed in post-mitotic cells throughout the dorsal-ventral axis of the spinal cord in *mib1^ta52b^* mutants (Fig. 7). As predicted from the *elavl3* results, for each of these genes, we see expression throughout most of the spinal cord, both medially and laterally (Fig. 8 G-L), although, consistent with the lateral view (Fig. 4P), the expression for *sall3a* (Fig. 8J) is less extensive than that of the other genes.

## Discussion and Conclusions

In this paper, while we, for completeness, include some expression data for other tissues, we primarily describe the expression of 21 different transcription factor genes and 1 transcriptional regulator gene in the zebrafish spinal cord. Given the high conservation of spinal cord development and transcription factor expression in vertebrates ^31,32^, these data are relevant not only to zebrafish development, but also to other vertebrates, including mammals and humans. All of these genes have broad, but not ubiquitious, expression patterns in the spinal cord. This suggests that they have specific functions in particular subsets of spinal cells, rather than general house-keeping roles in all cells. In order to hypothesize about the possible functions of these genes, we first need to know whether they are expressed in progenitor and/or post-mitotic cells in the spinal cord. At the stages that we examined, in WT embryos, spinal cord progenitor cells are located medially and as cells become post-mitotic, they move laterally. In contrast, spinal cord progenitor domain expression is lost (e.g. *dbx2*, Fig. 8A, B) and post-mitotic expression (e.g. *elavl3*, Fig. 8C, D) is usually expanded into additional cells in *mib1^ta52b^* mutants Based on our cross-sectional analyses and the dramatic reduction of their spinal cord expression in *mib1^ta52b^* mutants, we conclude that *aurkb, foxb1b, her8a, wdhd1*, and *zfhx3b* are all expressed in spinal cord progenitor cells (Figs. 2 and 7). *mybl2b, nr2f1b, sox19b*, *foxb1a, homeza*, *ivns1abpb, sall4, sox2*, *sp8b*, *znf1032* and *znf804a* are also predominantly expressed in progenitor cells, but *mybl2b, nr2f1b, sox19b*, *foxb1a, homeza* and *ivns1abpb* are also expressed in some dorsal post-mitotic cells, whereas *sall4, sox2*, *sp8b*, and *znf1032* are expressed in some post-mitotic cells in both the dorsal and ventral spinal cord and *znf804a* is expressed in only some ventral spinal cord post-mitotic cells (Figs. 3 and 7). In contrast, *myt1a, onecut1, sall1a, sall3a, sall3b*, and *tsc22d1* are predominantly expressed by different subsets of post-mitotic spinal cord cells, although our cross-sectional analyses suggest that *sall1a, sall3b* and *tsc22d1* are also expressed, at variable extents, in spinal cord progenitor cells (Figs. 4 and 7).

This is the first detailed description of spinal cord expression for most of these genes, in any vertebrate. The exceptions are *her8a* ^33,34^, *sall1a* ^35^, *sall3a* ^35^, *sox2* ^36–39^, *sox19b* ^37,40–42^ and *sp8b* ^43,44^. There is also some limited spinal expression data at 24 h for *aurkb* ^45,46^ ^supplementary^ ^data^, *foxb1b* ^47,48^, *nr2f1b* ^49^ ^supplementary^ ^data^, *sall4* ^50^, *zfhx3b* ^51^, as well as at stages earlier than those we examine in this paper for *foxb1a* ^47^, *foxb1b* ^52^, *myt1a* ^53^ and *sall1a* ^35^. Finally, there are some data at www.ZFIN.org or https://ibcs-bip-web1.ibcs.kit.edu/ffdb/ from large scale zebrafish expression screens, that show spinal cord expression of *aurkb*, *foxb1a, foxb1b, her8a, homeza, ivns1abpb, mybl2b, nr2f1b, onecut1, sall1a, sall3b, sall4, sox2, sox19b, sp8b, tsc22d1, wdhd1* and *zfhx3b* ^6–11^. For *homeza, ivns1abpb, mybl2b, onecut1, tsc22d1,* and *wdhd1,* these online database photos are the only spinal cord expression data that we are aware of, outside this paper. To our knowledge, there is no other expression data, in any tissue, in any vertebrate, for *znf1032* and no spinal cord expression data in any vertebrate for *znf804a*, not even in online databases of large-scale expression screens.

For all but one of the genes where there is some additional spinal cord expression data, our results are consistent with these other reports. The one possible exception is *sall3b*, where a photo from a large scale expression screen shows unrestricted expression, although no lateral views are provided so it is possible that this experiment just had higher levels of background expression than our data ^7^. However, for several genes, the data from the other sources shows weaker / less apparent spinal cord expression than our results and / or does not show spinal cord expression at stages older than 24 h. This is probably because many of these studies were not concentrating specifically on the spinal cord and, therefore, developed their *in situ* hybridization staining reactions to a level more appropriate for examining expression in other tissues. Spinal cord expression is often weaker than, for example, brain expression, particularly at later stages ^22,23^.

Notably, even in cases where some expression in the spinal cord has previously been shown, with the exception of *sall1a* ^35^, *sall3a* ^35^, *sox2* ^39,54^ and *sp8b* ^44^ there are no cross-sectional analyses of spinal expression and, with the exception of *her8a* ^55^, there is no analysis of spinal cord expression in *mib1* mutants, which means that it is often not clear whether the reported spinal cord expression is in progenitor and / or post-mitotic cells. This is a crucial piece of information for considering the functional roles of these genes in the spinal cord. In this paper, we have analyzed these aspects of expression for all 22 genes.

Interestingly, as described above, we find that while all of the genes that we examined are broadly expressed in the spinal cord, most of them have distinct expression patterns from one another. This suggests that they have different, specific functions in the spinal cord. However, for most of these genes, there is currently no data to suggest what their function(s) may be in the spinal cord. The exceptions are *aurkb,* which may be important for axonal outgrowth of spinal motor neurons ^56^, *her8a,* which is important for Notch signaling and, hence, neurogenesis e.g. ^34,55,57^, *onecut1,* which data from mouse and chick suggest may be important for motoneuron and V1 interneuron (Renshaw cell) development ^58–60^, *sox2,* which is required for correct differentiation of spinal motoneurons and oligodendrocytes ^39,61,62^ and *sp8b,* which is required for correct specification of the pMN/p3 progenitor domain boundary and hence the correct development of the cells that develop from these domains ^44,63^. Interestingly, at earlier stages of spinal cord development, *sall4* and *sp8b*, might belong to a common gene regulatory network as *sall4* is thought to regulate *pou5f3* expression, which in turn regulates *hoxb1a/b* expression in the posterior neurectoderm that forms the spinal cord, and *hoxb1a/b* over-expression can induce expression of *sp8b* ^64–67^. Further eludidation of these functions and the roles of the other genes in spinal cord development await further studies.

In conclusion, this study identifies 21 transcription factor genes and 1 transcriptional regulator gene with specific spinal cord expression patterns, that may have important roles in spinal cord development. In this way, it provides key new knowledge for Developmental Biologists, especially those interested in CNS development, and / or specific transcription factors, regulatory genes, or gene regulatory networks.

## Experimental Procedures

### Ethics statement

All zebrafish experiments in this research were carried out in accordance with the recommendations and approval of Syracuse University Institutional Animal Care and Use (IACUC) committee.

### Zebrafish husbandry and fish lines

Zebrafish (*Danio rerio*) were maintained on a 14-h light/10-h dark cycle at 28.5^◦^C. Embryos were obtained from natural paired and/or grouped spawnings of WT (AB, TL or AB/TL hybrid) fish, or heterozygous *mib1^ta52b^* mutants ^14^.

### *in situ* hybridization

We fixed embryos in 4% paraformaldehyde / phosphate-buffered saline (PBS) and performed single *in situ* hybridization experiments as previously described ^17,68^. For WT experiments, 30 embryos were processed per tube. For *mib1^ta52b^* experiments, 30 embryos from an incross of heterozygous *mib1^ta52b^* mutant parents were also processed per tube. Mendelian genetics predicts that we would, therefore, have about 7-8 mutant embryos per tube. For the experiments in this paper, we had a minimum of 6, and a maximum of 10 *mib1^ta52b^* mutant embryos per tube. Both mutant and sibling embryos were treated identically and stained for the exact same length of time to enable the comparison of expression patterns between genotypes. Since *in situ* hybridization is not quantitative, when comparing *mib* mutants and sibling embryos, we concentrated mainly on whether gene expression was expanded into additional cells or lost from specific domains. In some cases, we also saw stronger or weaker staining in the mutants. As the embryos were processed identically, this probably reflects changes in the level of expression, but to be sure of this, more quantitative methods, such as spatial transcriptomics, would need to be used, which are beyond the scope of this study.

For all genes except *dbx2*, we synthesized *in situ* RNA riboprobes using the methods described in ^18^ using the primers and annealing temperatures shown in Table 2. To avoid cross-reactivity, whenever possible, riboprobes were designed against 3’UTR or coding sequence lacking all conserved protein domains in Pfam ^69^. Primers were designed using Primer3 web version 4.1.0 at https://primer3.ut.ee ^70,71^ and the following design parameters: optimum primer size: 22 bp (minimum: 20 bp, maximum: 25 bp), optimum annealing temperature: 58.0°C (minimum: 57.0°C, maximum: 60.0°C), and optimum GC content: 50% (minimum: 40%, maximum: 60%). The preferred product size range was 800-1100 bp. This was not always possible, if there was little or no novel coding and/or 3’ UTR sequence available (see Table 2). The PCR conditions were: 98.0°C for 30 seconds, 35 cycles of: 98.0°C for 10 seconds; annealing temperature in Table 2 for 30 seconds and 72.0°C for 30 seconds, followed by a final extension for 5 minutes at 72.0°C. The PCR product was assessed on a 1% TAE gel, before purifying through phenol:chloroform:isoamyl alcohol extraction and precipitation with 0.2 M NaCl and ice-cold ethanol. If non-specific banding was generated in addition to the desired PCR product, the specific product was purified from the agarose gel using the Monarch DNA Gel Extraction Kit (NEB, T1020S). Each reverse primer contains the T3 RNA Polymerase minimal promoter sequence (shown in bold and underlined in Table 2). The template for the *dbx2 in situ* RNA riboprobe was prepared from plasmid DNA kindly provided by Gribble and colleagues ^26^. For all RNA riboprobes except *dbx2*, in *situ* probe synthesis was performed using 1 µg purified PCR product, T3 RNA Polymerase (11031171001, Roche) and DIG RNA Labeling Mix (11277073910, Roche). To synthesize the *dbx2* RNA riboprobe, 1 µg of linearized plasmid DNA was used, together with T7 RNA Polymerase (10881775001, Roche) and DIG RNA Labeling Mix (11277073910, Roche).

**Table 2.**
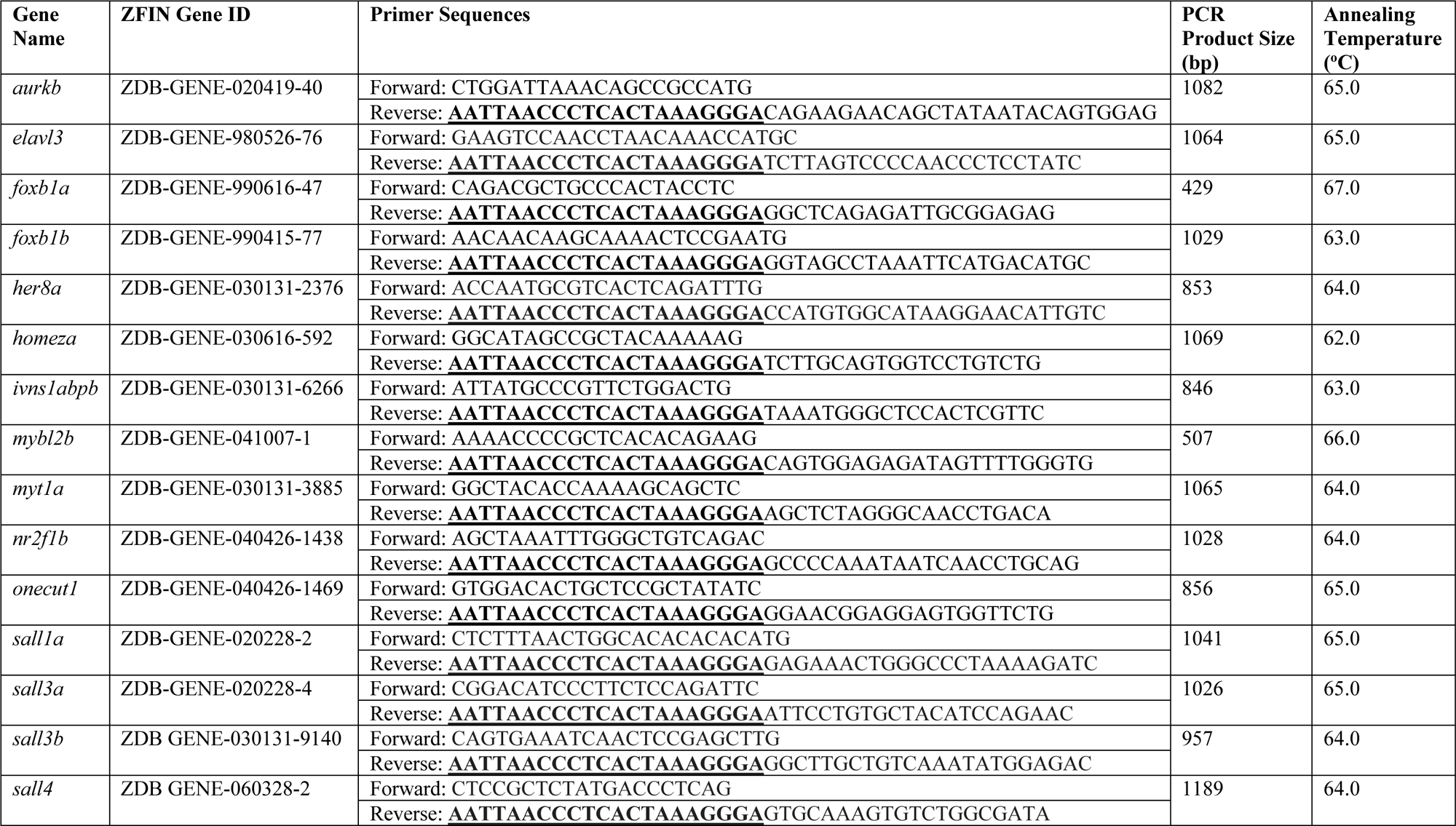
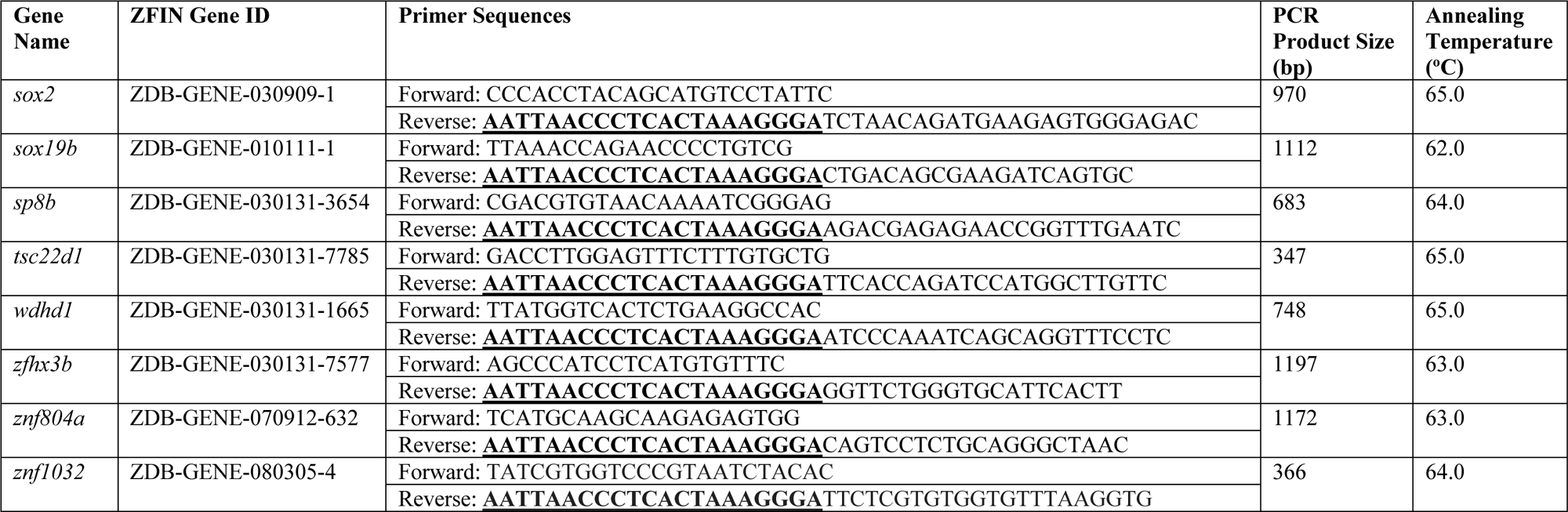
Gene Names, ZFIN Gene Identifiers, Primer Sequences and PCR Conditions for *in situ* Hybridization Riboprobe Synthesis. Column 1 lists genes analyzed in this study. Column 2 provides the unique ZFIN identification number for each gene. Columns 3 and 4 contain the primer sequences and expected product sizes (in base pairs (bp)) respectively, used to generate templates for anti-sense RNA riboprobe synthesis from 27 h WT cDNA. Bold and underlined text in column 3 shows the T3 RNA Polymerase minimal promoter sequence added to the reverse primers. Column 5 indicates the annealing temperature for the primer pairs shown in each row of column 3. For further conditions for riboprobe synthesis, please see Experimental Procedures.

Embryos older than 24 h were usually incubated in 0.003% 1-phenyl-2-thiourea (PTU) to prevent pigment formation. For some experiments, we added 5% of Dextran Sulfate to the hybridization buffer. In cases where this was done it is indicated in the relevant figure legend. Dextran sulfate can increase specific staining in *in situ* hybridization experiments as it facilitates molecular crowding ^72,73^.

In cases of low riboprobe hybridization efficiency, we exposed embryos to prolonged staining. In some cases, this produced higher background (diffuse, non-specific staining), especially in the hindbrain, where ventricles can sometimes trap anti-sense riboprobes ^23^.

### Imaging and Expression Analysis

Prior to imaging, all embryos for each gene were placed in a small dish of 70% glycerol / 30% sterile water, and examined using a stereo microscope (SMZ1000, Nikon), equipped with a gooseneck LED microscope light. We examined the expression pattern of each gene in each embryo to check if it was reproducible. If we observed variable staining within a genotype (for example, due to microclimates being formed during the *in situ* hybridization process which can cause differential exposure of the embryos to experimental reagents), we repeated the experiment. Once satisfied that we had reproducible staining, a minium of 5 embryos per gene, for each genotype, were analysed further as described below. *mib1^ta52b^* mutant embryos have an obvious morphological phenotype of curled tails and large heads and, in addition, the expression patterns of all of the genes that we examined were clearly changed in these mutants compared to their siblings. Therefore, we could not examine expression patterns blind to genotype.

All embryos were deyolked in 70% glycerol / 30% sterile water using mounting pins. For lateral and dorsal views of the embryo, whole embryos were mounted in 70% glycerol between coverslip sandwiches (24 mm x 60 mm coverslips; VWR, 48393-106), with 2-4 coverslips (22 mm x 22 mm; VWR, 16004-094) on either side of the sample to avoid sample compression. Cross-sections were cut by hand using a razor blade mounted in a 12 cm blade holder (World Precision Instruments, Cat. # 14134). Differential interference contrast (DIC) pictures were taken using an AxioCam MRc5 camera mounted on a Zeiss Axio Imager M1 compound microscope. All images were processed for brightness-contrast and colour balance using Adobe Photoshop software ^74^. Images of control and *mib1^ta52b^* mutant embryos are from the same experiment and they were processed identically. Figures were assembled using Adobe Photoshop ^74^.

## Author Contributions

S.J.E. designed the primers for all *in situ* hybridization RNA probes. S.J.E., P.C.C., and G.G. performed *in situ* hybridization RNA riboprobe synthesis. P.C.C. performed the majority of *in situ* hybridization experiments, with additional assistance from S.J.E., R.L.B., G.G., and W.F.F. R.L.B. performed most of the WT embryo cross-sectioning experiments, with additional help from S.J.E. S.B. performed all of the *MIB E3 ubiquitin protein ligase 1* embryo cross-sectioning experiments and imaging. S.J.E. performed the majority of the rest of the imaging, with assistance from P.C.C., R.L.B., S.B., and W.F.F. S.J.E. performed much of the data analysis and made all figures and tables in the paper. K.E.L. conceptualised and directed the study, acquired the financial support for the project, contributed to data analysis, and wrote the paper. All authors read and commented on drafts of the paper and approved the final version.

## Acknowledgements

We would like to thank Jessica Bouchard and several SU undergraduate fish husbandry workers for help with maintaining zebrafish lines and Richard Dorsky for the *dbx2* plasmid. This work was funded by NINDS R21NS073979, NINDS R01 NS077947 and NSF IOS 1755354 to K.E.L.

## Grant Sponsors and Numbers

NSF IOS 1755354, NINDS R21NS073979, NINDS R01 NS077947

